# Metabolism modulates network synchrony in the aging brain

**DOI:** 10.1101/2020.04.17.047233

**Authors:** Corey Weistuch, Lilianne R Mujica-Parodi, Anar Amgalan, Ken A Dill

## Abstract

Brain aging is associated with hypometabolism and associated global changes in functional connectivity. Using fMRI, we show that network synchrony, a collective property of brain activity, decreases with age. Applying quantitative methods from statistical physics, we provide a generative (Ising) model for these changes as a function of the average communication strength between brain regions. In particular, we find healthy brains to be poised at a critical point of this communication strength, enabling a balance between segregated (to functional domains) and integrated (between domains) patterns of synchrony. However, one characteristic of criticality is a high sensitivity to small changes. Thus, minute weakening of pairwise communication between regions, as seen in the aging brain, gives rise to qualitatively abrupt changes in synchrony. Finally, by experimentally modulating metabolic activity in younger adults, we show how metabolism alone–independent of other changes associated with aging–can provide a mechanism for global changes in synchrony.

## 1. Significance Statement

The brain is a biological machine that utilizes chemical energy to process information. However, the mechanism by which the brain adapts to resource constraints is poorly understood. This is particularly relevant in the aging brain, for which the ability of neurons to utilize their primary energy source, glucose, is diminished. Here, we provide a data-driven quantitative model for how brain-wide activity patterns are controlled by resource availability. This model shows that the brain is poised at a critical point, past which even minute changes in glucose utilization cause communication across the brain to markedly re-configure. Together, our results suggest that the clinical trajectory of cognitive changes associated with aging is discontinuous and can be mediated by metabolism.

## 2. Introduction

One of the most fundamental questions in neuroscience is how the familiar patterns of collective, brain-wide activity arise from the properties of the constituent neurons and their networks. Here, we study how the brain’s global activity patterns change with age, and how those changes might arise from the reduced metabolic activity of the constituent regions.

We draw on two types of experimental evidence. First, as established using positron emission tomography (PET), older brains show reduced glucose metabolism [1, 2, 3]. Second, as established by functional magnetic resonance imaging (fMRI), aging is associated with weakened *functional connectivity* (FC), *i.e.* reduced communication (on average) between brain regions [4, 5, 6]. Combining both observations suggests that impaired glucose metabolism may underlie changes in FC [1, 7]. Further supporting this link are studies showing disruptions similar to those seen with aging in Type 2 diabetic subjects [8, 9].

In healthy brains, resting-state brain activity (states during which subjects are not engaged in any explicit task) alternates between *segregating* computations to localized functional domains and *integrating* this information across these domains [7, 10, 11, 12, 13]. The metabolic cost of these activities increase in proportion to the number and length of functional connections between pairs of brain regions [14], making highly-connected (integrated) networks more energetically costly [10]. Moreover, connections with the highest cost are the first to weaken with age [6, 7, 15]. Thus, it has been hypothesized that declining glucose metabolism in older brains drives the loss of high-cost (integrated) functional activities [14]. Yet the relationship in aging brains, between energetic constraint at the level of individual brain regions and the apparent re-organization of the functional connectome, is still not well understood.

Here, we develop a generative model that describes how the probability distribution of FC patterns transforms with changes in global variables (such as age and metabolic activity)[16]. The approach of choice to understand how these changes arise is statistical physics, which interprets the collective properties of complex systems in terms of individual interactions between the underlying parts [17]. In particular, we employ an *Ising model* [18, 19, 20] to describe how pairwise interactions between brain regions give rise to specific profiles of network *synchrony*, a time-dependent average of the activity over the entire brain [21, 22, 23]

While the Ising model provides a general tool for describing the collective properties of complex systems, we adapt it to examine the specific relationship between brain aging and metabolic activity. To achieve this, we re-analyzed two fMRI datasets. The first is the *lifespan* Cam-CAN 3T fMRI resting state dataset of 636 individuals, ranging over ages 18-88 [24]. The second, in which we hold age constant in order to isolate the effects of metabolic activity alone, is the PAgB 7T fMRI within-subject experiment of 12 healthy young adults scanned while on glycolytic and ketogenic *diets* [25]. Ketone bodies decrease the relative free energy of ATP production by 27% as compared to glucose [26]. This additional efficiency of ketone bodies as a metabolite, observed even in healthy subjects, has been shown to increase both cardiac efficiency [26] as well as brain activity [25].

The significance of this work is three-fold. First, in contrast to the tools commonly used to study fMRI networks, our approach provides a predictive mechanism for how FC patterns change, in qualitatively significant ways, as a function of the average interaction between brain regions [16]. Second, we establish a direct link between network synchrony and the relative frequencies of integrated (high-cost) and segregated (low-cost) brain activities [10, 14]. Finally, we illustrate a precise relationship between differences in FC over the lifespan as well as in response to changes in the brain’s access to energy.

## 3. Methods

### 3.1. Lifespan and metabolic neuroimaging datasets

To identify how the collective features of fMRI change across the lifespan, we analyzed a large-scale 3T fMRI dataset: the Cambridge Centre for Ageing and Neuroscience stage II (Cam-CAN: ages 18-88, *N* = 636) [24]. The Cam-CAN study was designed to identify neural correlates of normal aging and provides a roughly uniform coverage of age groups, allowing comparison between groups as well as a wide array of behavioral measures. While the functional MRI imaging of Cam-CAN stage II included both task and resting state data, we used only resting state data, for which most regions of the brain have roughly similar statistical properties (see *Supplementary Fig. 2*). To relate these changes to energy in the brain, we additionally analyzed 7T fMRI data from the Protecting the Aging Brain (PAgB) database [25]. In a within-subjects experiment, young healthy adults (*N* = 12, *μ_age_* = 28± 6.73 years; 4 female) were scanned at resting state under two conditions: (1) *glycolytic*, following their standard diet, without fasting; and (2) *ketogenic*, following a high-fat, moderate-protein, low-carbohydrate (< 50 g/day) diet for one week, by which point all participants were in ketosis (> 0.6 mmol/L ketone blood concentration). For details on the glycolytic and ketogenic dietary regimes, as well as validation of their blood values and neurobiological effects as comparable to calorie-matched administration of glucose and D-*β*-hydroxybuterate, see previous work [25]. Studies were approved by the Institutional Review Boards of Cambridge University and Massachusetts General Hospital/Partners Healthcare, respectively; all participants provided informed consent.

### 3.2. MRI acquisition

The Cam-CAN lifespan dataset includes multiple imaging modalities (T1 and T2-weighted images, diffusion-weighted images, BOLD EPI images during tasks of three varying levels of cognitive demand, MEG images during two separate cognitive loads and magnetisation-transfer images). Of these, the resting state BOLD EPI fMRI was the focus of our analysis (full dataset documentation at [24]). The Cam-CAN functional imaging was done at 3T field strength over 8 min 40 s. The neuroimaging experiments of Cam-CAN study were conducted in Cambridge, UK at the Medical Research Council Cognition and Brain Sciences Unit (MRC-CBSU). Specifics of the BOLD EPI imaging protocol included: TR = 1970 ms, TE = 30 ms, flip angle = 78°, voxel size = 3 × 3 × 4.44 mm, slices = 32, number of measurements = 261. The PAgB metabolic dataset was acquired at ultra-high-field (7T) field strength at the Athinoula A. Martinos Center for Biomedical Imaging. Imaging included whole brain BOLD, field map, and T1-weighted structural (MEMPRAGE) images. BOLD images were acquired using a protocol quantitatively optimized, using a dynamic phantom, for detection-sensitivity to resting state networks [27]: SMS slice acceleration factor = 5, R = 2 acceleration in the primary phase encoding direction (48 reference lines) and online GRAPPA image reconstruction, TR = 802 ms, TE = 20 ms, flip angle = 33°, voxel size = 2 × 2 × 1.5 mm, slices = 85, number of measurements = 740 in each resting state session, for a total acquisition time of 10 minutes. Field map images were acquired using the following parameters: TR = 723 ms, TE1 = 4.60 ms, TE2 = 5.62 ms, flip angle = 36°, voxel size = 1.7 × 1.7 × 1.5 mm, slices = 89, for a total acquisition time of 3 min 14 s. The whole-brain T1-weighted structural volumes were acquired using a conventional multi-echo MPRAGE (MEMPRAGE) sequence with 1 mm isotropic voxel size and four echoes with the following protocol parameters: TE1 = 1.61 ms, TE2 = 3.47 ms, TE3 = 5.33 ms, TE4 = 7.19 ms, TR = 2530 ms, flip angle = 7°, with R = 2 acceleration in the primary phase encoding direction (32 reference lines) and online GRAPPA image reconstruction, for a total volume acquisition time of 6 min 3 s.

### 3.3. MRI pre-processing

Lifespan dataset pre-processing was conducted in the FMRIB Software Library (FSL; https://fsl.fmrib.ox.ac.uk/fsl/fslwiki/) and FreeSurfer (https://surfer.nmr.mgh.harvard.edu/): anatomical images were skull-stripped using FreeSurfer and co-registered to Montreal Neurological Institute (MNI) templates and mean functional images using FLIRT (part of FSL). Functional images were motion and fieldmap-corrected (using MCFLIRT and epidewarp), brain-extracted (using BET), and co-registered to MNI templates using transformations learned through the anatomical image. Motion parameter as well as tissue segmentation-extracted white-matter and CSF confounds (using FAST) were regressed out at ROI-level time series extraction stage using nilearn package (https://nilearn.github.io) [28]. Metabolic dataset pre-processing used Statistical Parametric Mapping 12 (SPM12; https://www.fil.ion.ucl.ac.uk/spm/software/spm12/) was used as in our previous studies conducted at the same acquisition parameters [29], [25]. Anatomical images (MEMPRAGE) were normalized to MNI templates using unified segmentation and registration. Images of each individual participant were realigned to account for head movements, and fieldmap-corrected (using epidewarp.fsl) for geometric distortions caused by the magnetic field inhomogeneity. Following normalization, structural images were probabilistically segmented into three tissues: grey matter, white matter, and cerebral spinal fluid. We did not apply spatial smoothing or global signal regression to pre-processing of either dataset. For all datasets, voxelwise data were parceled into the Willard 498 functional regions of interest (ROI) [30] corresponding entirely to grey matter voxels.

### 3.4. Ising model

Here we use the principle of maximum entropy [18, 20, 23] to build the minimally biased probability distribution of *N* binary (+1 or −1) node weights, 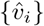 satisfying fixed constraints on the mean (0) and variance, *Var*(*s*) of the global property, 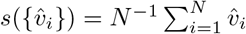 (synchrony). This is given by [23]:

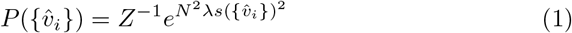

where *Z* is the partition function and normalizes the distribution. *λ* represents the average node-to-node interaction strength and is the basic mechanistic quantity of our model (**Fig. 1a, left**). Small values of *λ* describe networks in which interactions between nodes are weak and in which the node weights are independent of each other. In contrast, large values of *λ* describe networks in which interactions between nodes are strong and node activities are highly correlated (**Fig. 1a, right**). A given value of synchrony *s* may be obtained in many different ways; *i.e.,* it is *degenerate* (**Fig. 1b**). In other words, since there are 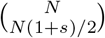 different ways to have 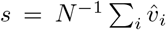, we find that the total probability *P*(*s*) of different synchronies is:

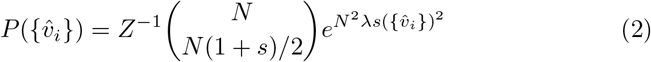

**Figure 1.**
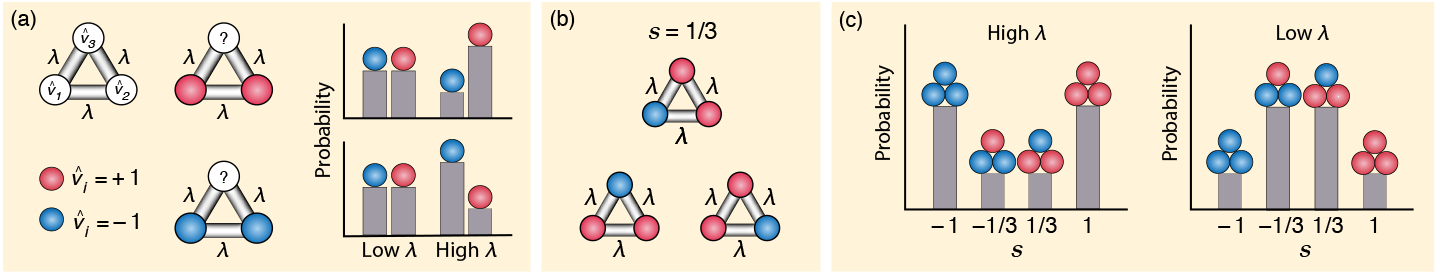
The Ising model predicts network probabilities from interactions between its nodes. (**a**) The Ising model maps binary variables onto a fully-connected network (**left**). Each variable (*i* = 1,2,… N) is a node with binary weight 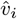 (represented by the colors red and blue), and each pair of nodes is connected by an edge with weight *λ*. Here we show the example of *N* = 3. The value of *λ* (> 0) describes the average interaction strength between nodes; the larger *λ* is, the more likely the unknown value of 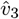 is to be similar to its neighbors (**right**). (**b,c**) The probability of each network is determined by its synchrony (s). (**b**) Multiple graphs give the same value of synchrony. Since there are 3 ways to have 1 blue node and 2 red nodes, there are 3 different graphs that give *s* = 1/3 (red minus blue divided by *N* = 3). This *degeneracy* effectively triples the probability of *s* = 1/3. (**c**) The probability distribution of *s* given by the Ising model is a function of *λ* and degeneracy. When *λ* (interaction) is large, the probability that |*s*| = 1 is large (**left**). But, when *λ* is small, degeneracy wins out and the probability that |*s*| = 1/3 is large (**right**).

Therefore, when *λ* is small, *P*(*s*) is determined by the degeneracy and low synchrony is most probable. Conversely, when *λ* is large, *P*(*s*) is determined by the interactions between nodes and high synchrony is most probable. In particular, as *λ* is varied, the relative importance of each of these terms changes. As can be seen in **Fig. 1c**, this causes *P*(*s*) to change from a bimodal (left) to a unimodal (right) distribution. The critical point, *λ_c_*, is the value of *λ* where this shift happens (i.e. when these two contributions are balanced). Using the standard approximation of the binomial coefficients, *P*(*s*) becomes:

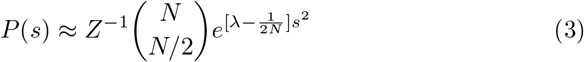

Conceptually, when *λ* < *λ_c_*, P(s) opens downwards like a Gaussian; *s* = 0 is most probable. However, when *λ* > *λ_c_*, *P*(*s*) opens upwards and large values of *s* (both positive and negative) are probable. When *N* = 498 (the number of regions), we find that this critical point is 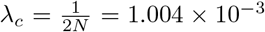, coinciding with the observed transition between unimodal and bimodal synchrony (**Fig. 1c**). To simplify our analysis, now refer to the rescaled interaction Λ: Λ = (*λ − λ_c_*)*/λ_c_*.

### 3.5. fMRI binarization

In order to access the time-dependent network properties of our data, we first binarize the fMRI time series. This method simplifies time series while preserving their functional connectivity (FC) patterns. In particular, the Pearson correlation *ρ*(*X, Y*) is widely used to estimate FC between arbitrary pairs of variables (*X*,*Y*):

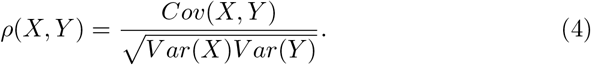

Here variables (*X* and *Y* for example) are the nodes of a graph and *ρ* is the weight of the edge between them. However, these connection strengths often change over time [31]. Thus, we calculate *ρ* over each pair of successive time points, reducing Eq. 4 to:

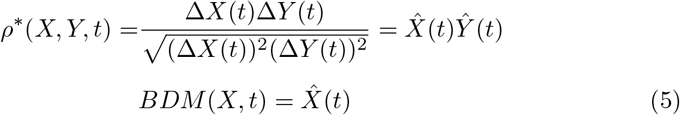

where 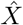 and *Ŷ* are the signs of the time derivatives of *X* and *Y* respectively and the time-dependent correlation, *ρ*^*^, is their product. This procedure takes our original time series *X*(*t*) and produces a simplified, binarized time series 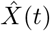 (*Binarized Derivative Method*, BDM). By computing these binarized values for long periods of time, we can ask questions about how the probabilities of different sequences (in time) and patterns (over regions) change with different conditions (such as with age and diet). As validation of this method, we find that this simplified representation preserves fMRI FC patterns across time (*Supplementary Fig. 1a*) and for different subjects (*Supplementary Fig. 1b*). This approach has two key advantages over previous methods [31, 32]. First, it simplifies complex, many-variable interactions in terms of dynamical patterns of binary (+1 and −1) variables. Second, it is naturally compatible with Ising-like models, which have been shown to be powerful tools in isolating latent relationships within networks of neurons [20, 23].

### 3.6. Model fitting

We then fit the Ising model to our data (**Fig. 2**). First, we took the fMRI signal *υ_i_*(*t*) for each region *i* and time *t* and binarized it using BDM. The model assumes that all nodes have, on average, similar FC strengths. We tested this assumption by computing the total (over all pairs) FC for each region, and we used the subject-averaged (over all diets and ages) FC matrix as our reference (*Supplementary Fig. 2*). From these signals, we found that most nodes are primarily positively correlated, while a few nodes were primarily negatively correlated with other nodes. For the latter, we flipped 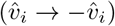 for these regions only in order to satisfy the assumptions of our Ising model (*Supplementary Fig. 2*). For each subject, we then computed the time-dependent *synchrony s*(*t*) (each TR is a time point) using the binarized fMRI signals from all (498) regions of the brain. We then took the histogram of *s*(*t*) for each subject to get a distribution *P*(*s*), giving the variation in synchrony per individual. This was then used to obtain Λ by fitting *P*(*s*) to the Ising model Eq. 2. This fit is expressed by the Bayesian posterior distribution *P*(Λ|Data), which captures the relative quality of of our model. We use a uniform (unbiased) prior distribution of Λ; thus the posterior is computed directly from the likelihood function 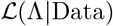 of our Ising model **Eq. 2**. In practice, we will summarize this posterior by its peak (the maximum likelihood estimate) and its width (error bars). As fMRI signals are auto-correlated, the data (*s*(*t*)) are not fully independent. To compensate for this effect, we consider conservative (0.01 likelihood ratio) error bars for Λ.

**Figure 2.**
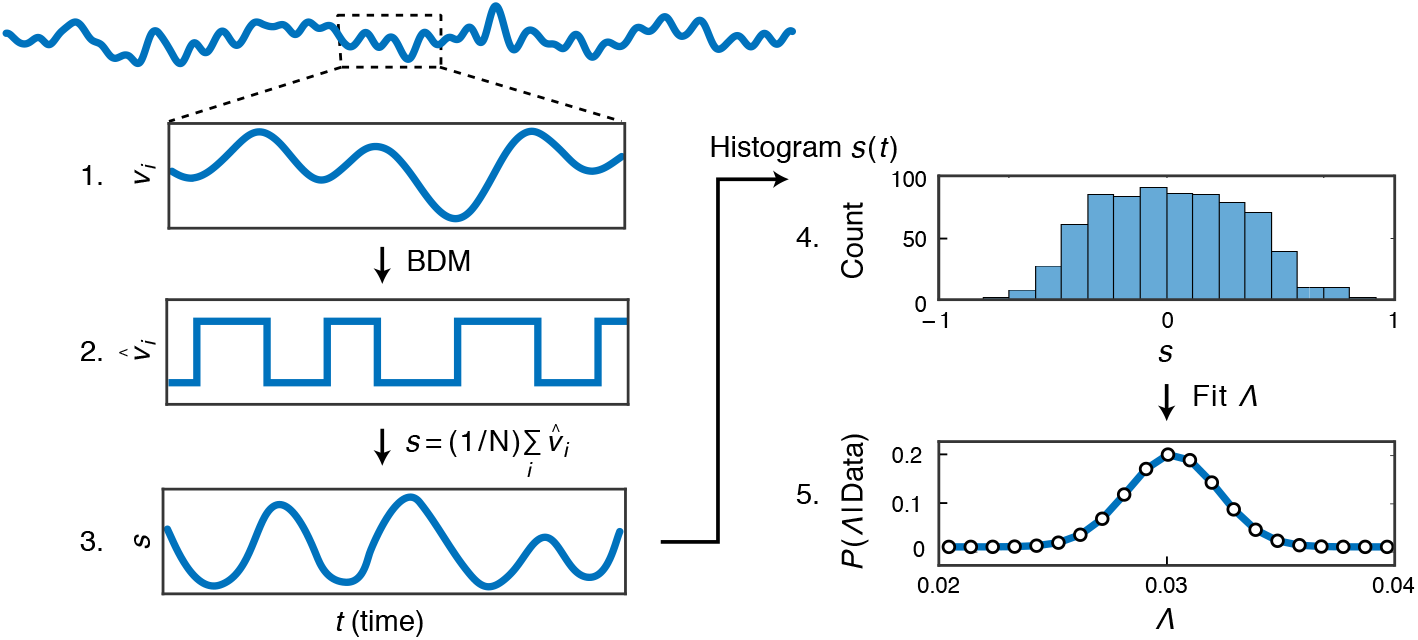
How we obtain the Ising model parameter Λ from fMRI data. [1] shows the fMRI signal *υ_i_*(*t*) from the *i*th brain region (out of 498), as a function of time *t*. [2] We binarize it, to give 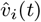. [3] The binarized signals are then averaged over all brain regions, giving that individual’s time-dependent *synchrony s*(*t*). [4] We then histogram into *P*(*s*) the different *s* values over time [4], to express the variations in an individual’s synchrony levels. [5] We then find the value of Λ that best fits *P*(*s*) for each individual. *P*(Λ Data) expresses the Bayesian posterior probability (with a uniform prior distribution over Λ) that our data *P*(*s*) was generated from an Ising model (**Eq. 2**) with relative interaction strength Λ.

## 4. Results

To interpret our fMRI data, we developed a generative biophysical approach based on a network *Ising model* [19, 20]. Widely used in physics, the Ising model describes how pairwise interactions among microscopic, binary (±1) elements give rise to macroscopic behaviors, including correlations ([19], **Fig. 1**). In other words, the Ising model allows us to describe time-dependent variability (probabilities) of different brain states for each subject.

We are particularly interested in the collective (i.e. regionally-averaged) properties of brain activity. In general, collective properties can often be described using mean-field models, where every component of interest is approximated as being connected to every other component with the same strength [23, 33]. Here the collective property of interest is the observed network *synchrony*, *s*, or the average activity across the 498 Willard Atlas brain regions measured in fMRI experiments [30, 23]. The probability distribution of different synchronies can then be described by a mean-field Ising model, with a single average interaction strength (assumed positive) between all pairs of brain regions (see *Supplementary Figs. 2 and 3* for further justification). To explore this model, we find the value of the interaction strength, Λ, that best fits the experimentally observed synchrony values for each subject (**Fig. 2**). Thus, each value of *s* corresponds to the degree of consensus of a particular network produced by the best-fitting Ising model [23]. As further validation for our model, we find that the Ising model, regardless of age and diet, correctly captures the kurtosis of *P*(*s*), a higher-order feature that cannot be generally predicted from correlations alone (*Supplementary Fig. 3*).

Ising models are useful in understanding how changes in smaller-scale properties (such as the interactions between brain regions) can give rise to abrupt and qualitatively distinct collective phenomena at larger scales. Much like water at its boiling point, which discontinuously changes from liquid to vapor, these changes occur at an intermediate value of the interaction strength, called the *critical point*. Here we use Λ to denote the deviation from the critical interaction strength (Λ = 0) of the Ising model. Figure 3 illustrates how the distribution of synchronies (with example brain networks shown for comparison) changes as a function of Λ, from unimodal (low synchrony, *s* = 0, blue) when Λ < 0 to bimodal (high synchrony, *s* = ±1, orange) when Λ > 0. While both low and high synchrony networks are equally likely at the critical point (**Fig. 3**, red, Λ = 0), small changes in Λ lead to large, abrupt changes in this balance.

**Figure 3.**
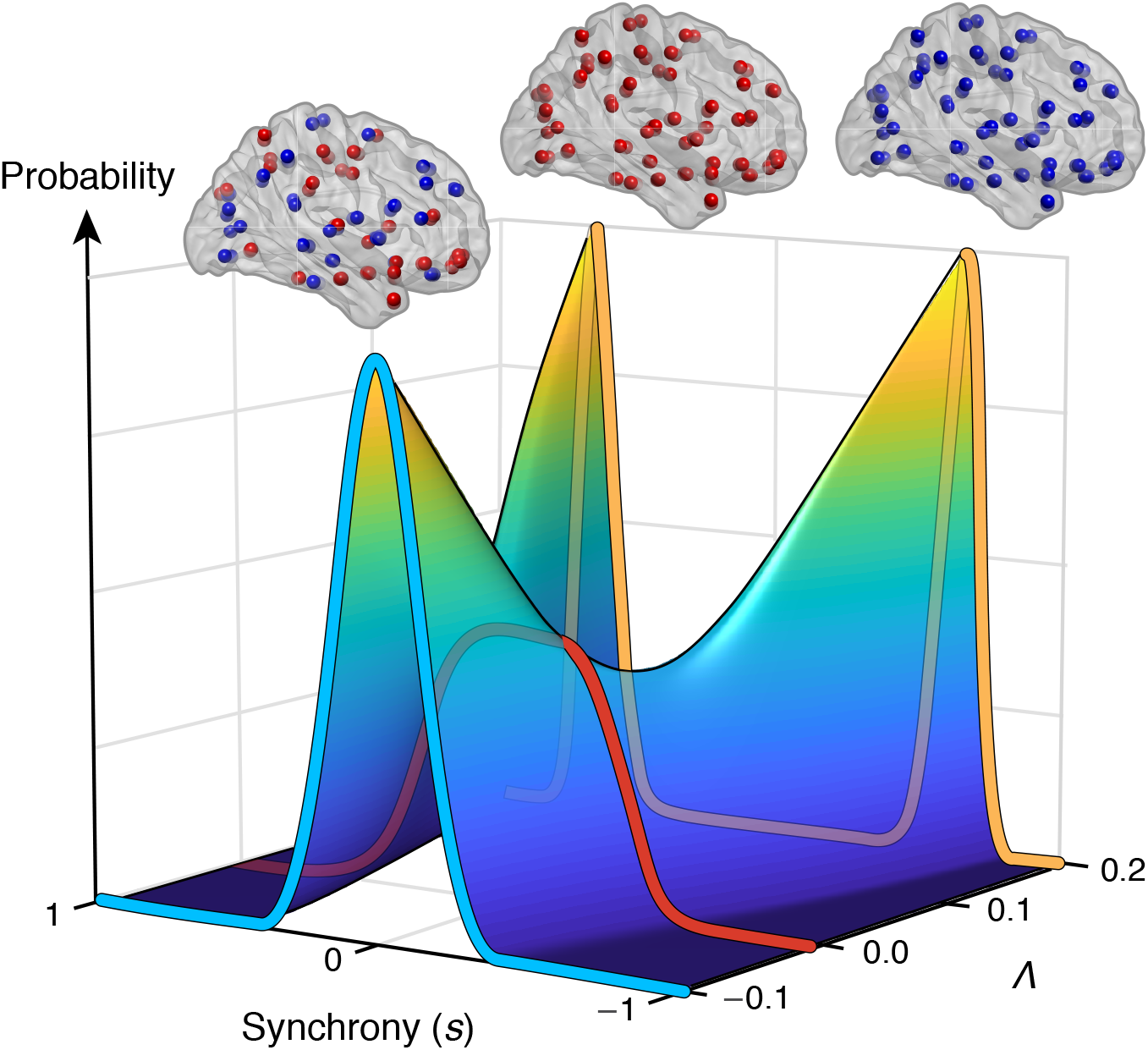
The Ising model applied to brain synchrony. Shown is the probability distribution of different values of synchrony (s) for different values of the dimensionless quantity Λ, reflecting the distance of the actual interaction strength, *λ* from the critical point *λ_c_*: Λ = (*λ* − *λc*)/*λ_c_*. For Λ < 0 (weak interactions), there is a single unimodal population having a peak at *s* = 0 (blue line). For Λ > 0 (strong interactions), the population is bimodal, with a peaks at *s* ≫ 0 and *s* ≪ 0 (orange line). Above each peak is an example network; nodes are brain regions and colors are states (red +1, blue −1). Λ = 0 defines the critical point, where *s* = 0 changes from a minimum to a maximum and P(s) rapidly changes (red line). At the critical point, low and high synchrony networks are equally probable.

To establish the relationship between synchrony and the occupation probabilities of specific functional networks, we separately computed the inter-subject average FC matrices during periods of low and high synchrony. During periods of low synchrony, functional connections are found to be sparse, favoring connections between local (segregated) networks of regions (**Fig. 4a**, Seg). In contrast, high synchrony networks are typified by dense connections (integrated) between multiple functional domains across the brain (**Fig. 4b**, Int) [10]. Consequently, just as with synchrony (**Fig. 3**), different values of Λ change the relative time spent in segregated (*P_Seg_*) and integrated (*P_Int_*) networks (**Fig. 4c**, *R*^2^ > 0.9, sigmoidal fit not shown, each colored marker is a subject), independent of age or diet. The time spent in each pattern was computed as the similarity of each subject’s FC to the extracted patterns, Int and Seg. When Λ < 0, low synchronies (i.e. segregated networks) occur more frequently, while the opposite holds when Λ > 0. In both cases, this balance rapidly shifts at the critical point, Λ = 0.

**Figure 4.**
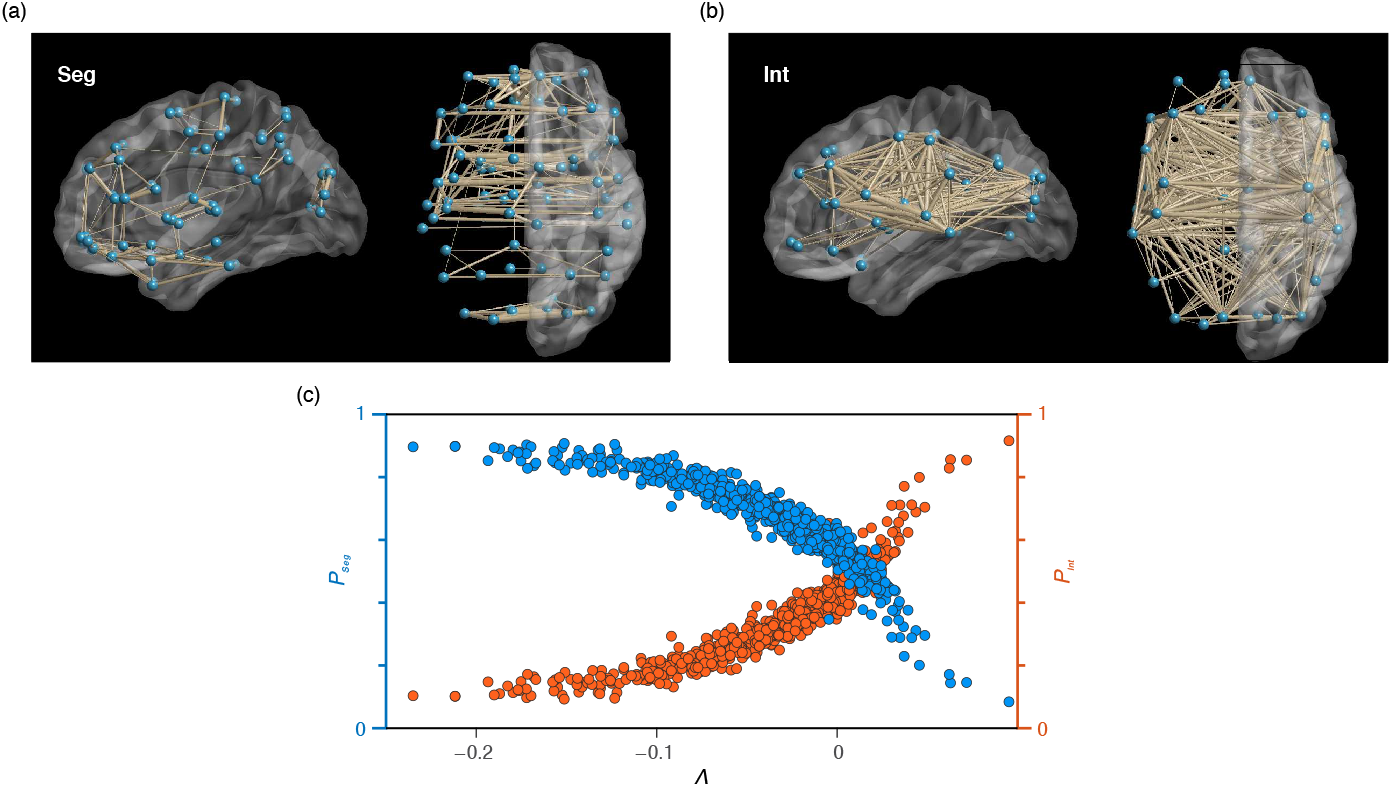
Λ controls the balance between segregated (low s) and integrated (high s) networks. (**a,b**) Inter-subject average functional connectivity (Pearson Correlation) during low (**a**) and high (**b**) synchrony, visualized using the BrainNet Viewer showing the top 10 % of connections [34]. (**a**) Low synchrony (*s* = 0) reflects segregation (Seg). (**b**) High synchrony (|*s*| > 1/2) reflects integration (Int). (**c**) The fraction of time each subject (each data point and their specific value of Λ) spends in integrated (*P_Int_*, orange) and segregated (*P_Seg_*, blue) networks. Time spent was calculated from a bivariate regression of the functional connectivity (here over all *s*, from each subject) with the patterns, Seg and Int. Λ < 0 corresponds to large *P_Seg_* and small *P_Int_* while Λ > 0 corresponds to the opposite. The cross-over in (**c**) occurs at the critical point, Λ = 0.

Changes in FC with both age and diet can be described by changes in the region-region interaction strength Λ. In particular, we find that Λ significantly decreases with age (*p* = 1.7 × 10^−38^, *N* = 636, **Fig. 5a**), suggesting that aging is associated with a marked shift from integrated towards more segregated network activities. But, upon switching from a lower-energy glycolytic to a higher-energy ketogenic diet, Λ increases (*p* = 1.2 × 10^−3^, *N* = 12, **Fig. 5b**) by about 25% to 50% of the decrease seen over the entire lifespan. Thus, by toggling the relative frequencies of segregated and integrated networks, Λ reflects an average cost of functional activity and, as suggested by our metabolic experiment, the amount of energy available to the brain. Thus one way the brain may conserve energy when this amount of energy available is decreased, such as through aging, is by decreasing Λ.

**Figure 5.**
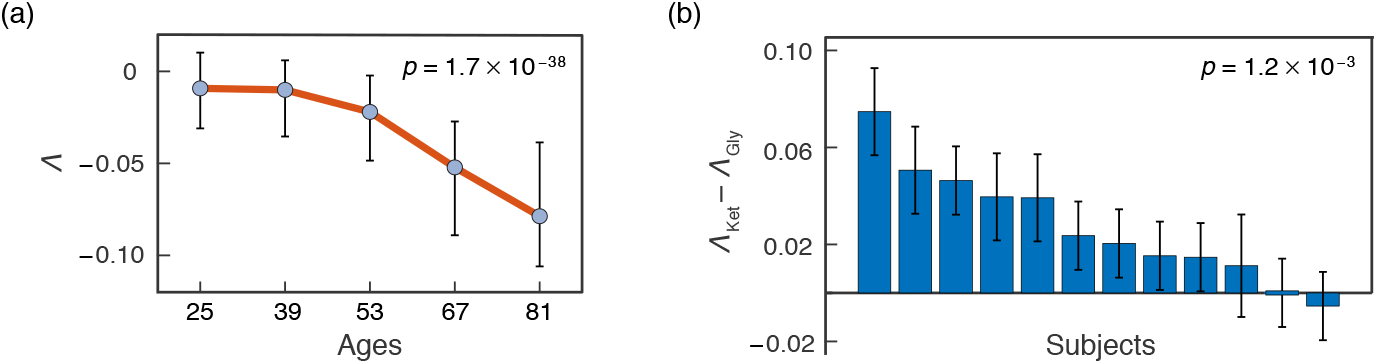
Λ significantly decreases with age (**a**, *p* = 1.7 × 10^−38^) and increases on the higher-energy, ketogenic diet (**b**, *p* = 1.2 × 10^−3^). (**a**) Each point (as well as the orange curve connecting them) reflects the median best-fit Λ values for each (of 5) equal width (14 years) age groups. Error bars represent the upper and lower quartiles. We used a Spearman-rank Permutation test (*N* = 636, *ρ*(634) = −0.48) to test significance of the nonlinear relationship between Λ and age. (**b**) Change in Λ for each subject (*N* = 12, *W* = 3) when switching from a lower-energy glucose (glycolytic, Gly) to higher-energy ketone (Ket) metabolism. Error bars reflect a 0.01 likelihood ratio confidence interval. A Wilcoxon 1-sided signed rank test (*N* = 12) was used to test if ketones significantly increased Λ.

But why would a small change in Λ lead to the dramatic changes in FC seen in older age? Precisely because young healthy brains are poised at the critical point (Λ = 0), very small changes in the interaction strength between regions lead to a sharp transition in the ratio of integrated to segregated networks [19, 20]. **Figure 6** expresses this in terms of the probability distribution of *s*, now viewed from the top-down. Here younger brains (green, age 25 ± 7) are near the critical point (black), allowing them to access both high and low synchrony networks. But as Λ (a proxy for energy availability, [14]) decreases, such as observed in older brains (yellow, age 81 ± 7), the probabilities of higher synchrony networks quickly fall to 0.

**Figure 6.**
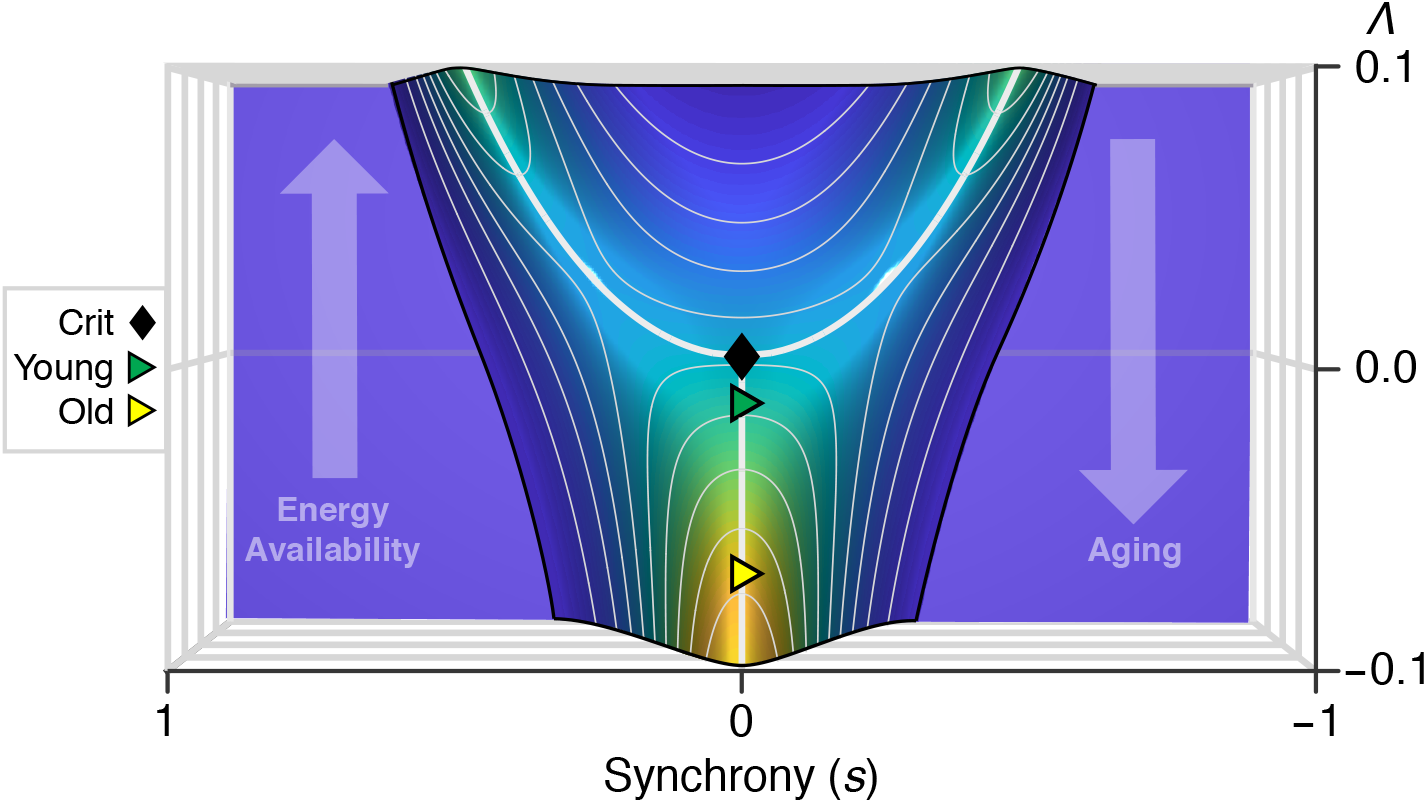
Younger brains are poised at a critical point; this is disrupted by decreasing energy availability. Shown is the probability distribution of synchrony (*s*) vs Λ, viewed from the top-down. At the critical point (Λ = 0, black), peak synchrony (indicated by a white line) changes from low (*s* = 0) towards high (*s* < 0 and *s* > 0) values. Near this transition, such as seen in younger brains (Age 25 ± 7, *N* = 85, green), both low and high synchrony networks can be accessed. Reducing energy availability causes Λ (through associated decreases in FC, [14]) to decrease. Older brains (Age 81 ± 7, *N* = 121, yellow) have smaller Λ and only access low synchrony networks. The plotted triangles correspond to the Λ values centered at ages 25 (Young) and 81 (Old) (**Fig. 5**).

## 5. Discussion

Our results suggest that the principal functional changes associated with aging, in terms of network synchrony, are controlled by an average interaction strength (Λ) between pairs of brain regions. Crucially, unlike graph theoretic features normally used to describe such data[16], Λ encodes *how* the aging brain rewires. We have also shown that Λ governs a trade-off between low-cost, segregated and high-cost, integrated activity patterns. Furthermore, as suggested by our findings, we hypothesize that Λ is decreased in older brains to compensate for glucose hypometabolism. But, because younger brains are poised near a critical point, this compensation results in sharp changes in functional connectivity.

It is important to note that aging and ketosis each exerts independent systemic effects that need to be considered in interpreting the results. For example, older subjects often have cardiovascular changes that affect neurovascular coupling[35] and thus, by extension, the blood oxygen level dependent (BOLD) response measured by fMRI. Likewise, ketosis has systemic effects, such as diuresis and thus lowered blood pressure, as well as reduced cellular need for oxygen, all of which also could theoretically affect BOLD. However, there are several reasons to suspect that these alternative mechanisms are not the sole causal influence of shifts in Λ. First, to minimize the primary cause of neurovascular confounds, the lifespan dataset specifically excluded individuals with cardiovascular disease, including cerebral ischeaemia [36]. Moreover, while the impact of arteriosclerosis in reducing the dynamic range of BOLD could reduce signal/noise and therefore reduce the strength of measured connections overall, it would not discriminate between integrated versus segregated networks and the transitions between them. Second, shifts in *λ* were observed not only in the aging dataset, but also in the dietary dataset, the latter of which included only younger individuals and thus eliminated systemic aging effects as a variable. Third, systemic (non-metabolic) effects of ketosis, such as reduced cerebral blood pressure and reduced need for oxygen, should decrease BOLD activation, while we have previously shown ketosis to increase BOLD activation, both in our dietary dataset as well as an independent dataset in which ketosis was achieved by administering exogenous D-*β*-hydroxybuterate[25]. Nevertheless, dissociating metabolic from more systemic influences of aging and ketosis is one important direction for our future research.

The metabolic cost of connectivity is known to reflect both signaling along axons as well as between synapses. As such, Λ may reflect the average synaptic connectivity across the brain, as suggested by recent evidence linking global resting state fMRI fluctuations to synaptic activity [21]. Indeed, synaptic connections weaken with age [37, 38] and are particularly vulnerable to metabolic disruptions [39, 40, 41, 42]. However, the fact that age was associated with a reduced probability of integrated activities (with longer connections) in favor of segregated activities (with shorter connections) suggests that the metabolic cost of axon conductance may also play a key role. Long-range connections are known to be disproportionately diminished not only with age [15] but also epilepsy [43], the latter of which commonly shows improvement with ketosis.

That brains at their presumed peak of functionality should be poised so close to a critical point of synchrony may reflect an evolutionary selective advantage. Criticality is not only a widely-observed feature of neural activity[44, 45, 46], but also enables the broadest range of functional patterns while also achieving maximum sensitivity to external drivers (e.g. sensory stimuli) [19, 47]. Some recent work suggests that signatures resembling criticality may be generic features of systems with many unobserved variables [48]. However, if this were the case, one would find these signatures in both younger and older brains, which is not consistent with our findings.

In conclusion, the Ising model provides a data-driven generative model for how the brain adapts to resource constraints, such as progressive glucose hypometabolism in aging brains. By simply shifting the balance between integration and segregation away from the critical point, the brain is able to modulate its fuel efficiency without the need to invest in new synaptic connections [7, 14]. Thus toggling Λ reflects an optimal strategy for the brain, enabling the smoothest adaptation for the smallest energetic cost. At the same time, the brain’s protective strategy in conserving energy may produce discontinuous trajectories for cognitive changes associated with aging, both in terms of diminished sensitivity to sensory stimuli (as predicted by shifts from criticality) as well as cognitive processing associated with flexibility in switching between both segregated and integrated networks.

## Data and code availability

Lifespan fMRI data are publicly available from Cam-CAN [24]. Metabolic fMRI data are located at Data Archive for the Brain Initiative (DABI: https://dabi.loni.usc.edu/explore/project/42) in the Protecting the Aging Brain (PAgB), Project 1926781 repository. Additional details (including links to custom MATLAB and Python codes used in the processing and analyses of data) can be found at http://www.lcneuro.org/software-and-instrumentation.

## Acknowledgments and Funding

The research was funded by the WM Keck Foundation (LRMP, KD), the NSF BRAIN Initiative (LRMP, KD: ECCS1533257, NCS-FR 1926781), and the Stony Brook University Laufer Center for Physical and Quantitative Biology (KD).

## CRediT authorship contribution statement

LRMP designed the experiments. AA pre-processed the data. CW did the modeling and designed the methods used. CW, LRMP, and KD wrote and revised the paper.

