## Supplementary Information for "Metabolism modulates network synchrony in the aging brain"

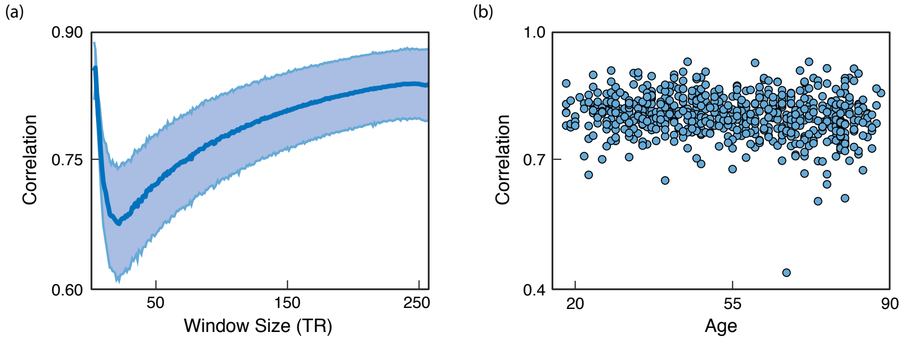


**Supplementary Figure 1** Functional Connectivity (FC) can be recovered from simple, binarized signals (BDM). (**a**) The correlation between FC calculated using raw and binarized data across different windows of time. Error bars represent the population standard deviation ($N = 636$). (**b**) The same correlations (using the maximum window size) plotted against age.

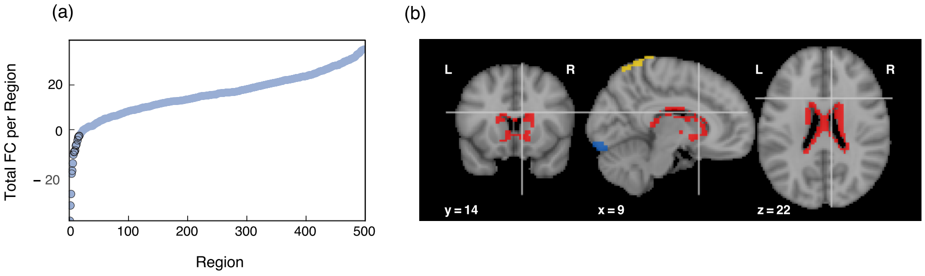


**Supplementary Figure 2** Resting-state functional connectivity (FC) separates the brain into two clusters: positive and negative. (**a**) The population-averaged ($N$ = 636 subjects) total functional connectivity of each region (498 in total, summed over the remaining 497 pairs) ordered from smallest to largest. The 16 leftmost regions (black highlighted circles) form the negative cluster, while the remaining majority form the positive cluster. (**b**) A three plane view of the negative (colored) regions. The negative cluster included various subcortical regions (thalamus, the fornix of the hippocampus, and the caudate nucleus in red), the supplementary motor area (yellow) and the lingual gyrus (blue). Our pre-processing excluded all non-grey matter voxels for each subject. Thus, while these regions are near grey matter borders, they were not an artifact of pre-processing.

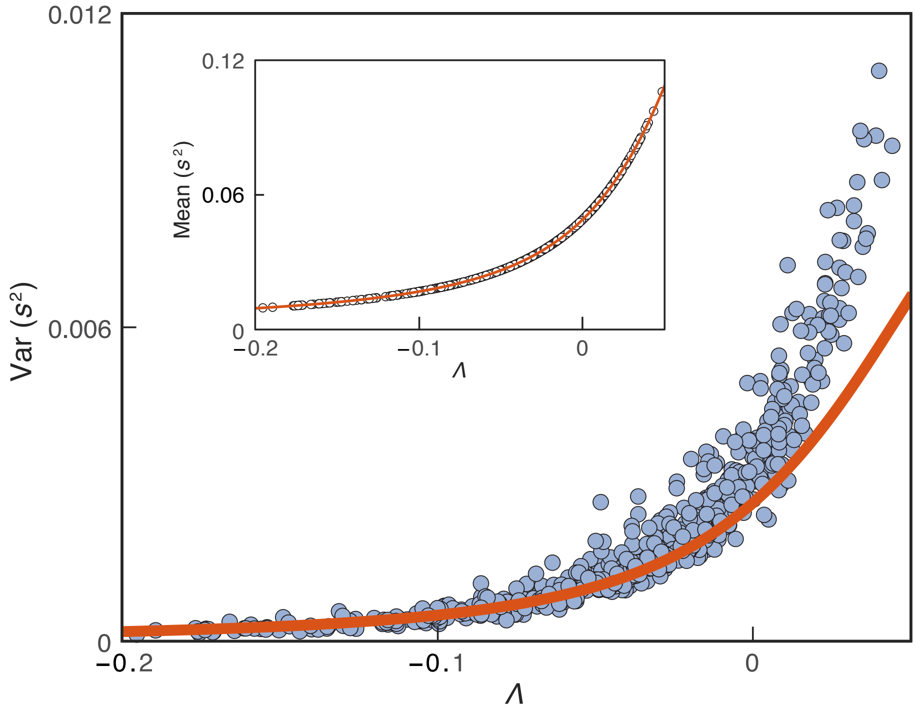


**Supplementary Figure 3** The Ising model predicts higher-order features of functional data. We fit binarized functional data from our lifespan and metabolic datasets to the model using Maximum Likelihood estimation. Shown are fourth moments of synchrony $Var(s^2)$, as predicted by the model (orange), compared to experimental data (each marker is a subject). In addition to the second moment Mean($s^2$), which is perfectly fit by design (inset), the model captures well the value of this additional feature sufficiently far up to $\Lambda > 0$ to be relevant to all the biological data we study here. Additionally, these deviations do not affect the important quantitative features of our model [1, 2].

[1] Sander Dommers, Cristian Giardinà, Claudio Giberti, Remco van der Hofstad, and Maria Luisa Prioriello. Ising critical behavior of inhomogeneous curie-weiss models and annealed random graphs. *Communications in Mathematical Physics*, 348(1):221–263, 2016.

[2] Ginestra Bianconi. Mean field solution of the ising model on a barabási–albert network. *Physics Letters A*, 303(2-3):166–168, 2002.
